# Novel *Plasmodium falciparum* metabolic network reconstruction identifies shifts associated with clinical antimalarial resistance

**DOI:** 10.1101/119941

**Authors:** Maureen A. Carey, Jason A. Papin, Jennifer L. Guler

## Abstract

**BACKGROUND:** Malaria remains a major public health burden and resistance has emerged to every antimalarial on the market, including the frontline drug artemisinin. Our limited understanding of *Plasmodium* biology hinders the elucidation of resistance mechanisms. In this regard, systems biology approaches can facilitate the integration of existing experimental knowledge and further understanding of these mechanisms.

**RESULTS:** Here, we developed a novel genome-scale metabolic network reconstruction, iPfal17, of the asexual blood-stage *P. falciparum* parasite to expand our understanding of metabolic changes that support resistance. We identified 11 metabolic tasks to evaluate iPfal17 performance. Flux balance analysis and simulation of gene knockouts and enzyme inhibition predict candidate drug targets unique to resistant parasites. Moreover, integration of clinical parasite transcriptomes into the iPfal17 reconstruction reveals patterns associated with antimalarial resistance. These results predict that artemisinin sensitive and resistant parasites differentially utilize scavenging and biosynthetic pathways for multiple essential metabolites including folate and polyamines, and others within the mitochondria. Our findings are consistent with experimental literature, while generating novel hypotheses about artemisinin resistance and parasite biology. We detect evidence that resistance parasites maintain greater metabolic flexibility, perhaps representing an incomplete transition to the metabolic state most appropriate for nutrient-rich blood.

**CONCLUSION:** Using this systems biology approach, we identify metabolic shifts that arise with or in support of the resistant phenotype. This perspective allows us to more productively analyze and interpret clinical expression data for the identification of candidate drug targets for the treatment of resistant parasites.

## BACKGROUND

Three billion people are at risk for malaria infection globally and treatment approaches are failing. Malaria is caused by *Plasmodium* parasites, and most deaths are associated with human-infective *P. falciparum*. Without an efficacious vaccine, antimalarials are essential to combat the severity and spread of disease. Combination therapies are implemented to preserve antimalarial efficacy and slow resistance development [1-3]; despite this approach, this eukaryotic pathogen has developed resistance to every antimalarial on the market [4-6].

Typically, resistance is conferred by genomic changes that lead to drug export or impaired drug binding (for example [7]); however, non-genetic mechanisms have also been implicated in *Plasmodium* resistance development [8-10] and other pathogenic organisms, such as *Pseudomonas aeruginosa* [11] (reviewed in [12]). These laboratory-based studies provide insight into metabolic flexibility but the presence of relatively few examples limit our understanding of this method of adaptation, especially in malaria. Here, we aim to look beyond genetic mechanisms of resistance to identify resistance-associated metabolic adaptation. We hypothesize that metabolic changes must occur to support the resistance phenotype and resistance-conferring mutations. Ultimately, these changes, or ‘shifts,’ are required to increase the fitness of resistant parasites, or support the development of additional genetic changes that affect fitness. Metabolic or phenotypic ‘background’ could be as important as genetic background in the development of resistance.

In clinical malaria infections, artemisinin resistance is established in Southeast Asia [13-15]. This phenotype is correlated with mutations in the *P. falciparum Kelch13* gene [13, 14, 16, 17] and changes in both signalling pathways [18-21] and organellar function [22-29]. Overall, due to the complexity of artemisinin’s mechanism of killing (see citations above and [22, 28, 30-36]), it has been challenging to separate the causes and effects of resistance. For this reason, there are few novel solutions to antimalarial resistance beyond altering the components of combination therapies to regain efficacy (e.g. artemisinin-atovaquone-proguanil [1]). We aim to gain a new perspective on resistance by viewing it through a ‘metabolic lens’. By characterizing the metabolic shifts that occur during or after resistance acquisition, we can begin to understand more about what it takes to support new functions, such as novel signalling (e.g. PI3K signalling is affected by *PfKelch13* mutations [14, 15, 20, 37, 38]), drug detoxification (e.g. regulating ROS stress associated with artemisinin treatment [24, 25, 30, 33]), or stage alterations (e.g. dormancy of early ring stages [18, 39-42]) in resistant parasites. Once we identify these compensatory changes, we can potentially target them. *Plasmodium* metabolic genes are better characterized than signalling pathways, as (for example) PlasmoDB identifies 43 3D7 genes associated with the term ‘signalling’ as opposed to 1112 3D7 genes associated with the term ‘metabolism’ [43], and many antimalarials target metabolic functions [44-47]. Moreover, metabolism has been described as the best-understood cellular processes [48], making interpreting metabolic analyses more tractable. Ultimately, if we can identify targetable conserved metabolic differences that arise with or in support of resistance, we can develop more robust antimalarial combination therapies aimed at preventing resistance.

Here, we use a systems biology approach to analyze the metabolic profile associated with resistant and sensitive parasites. First, to maximize the accuracy of our predictions, we curated an existing genome-scale network reconstruction of asexual blood-stage *P. falciparum* metabolism. Using constraint-based metabolic modeling, we integrated transcriptomic data from over 300 clinical isolates from Cambodia and Vietnam with varying levels of artemisinin sensitivity. This approach identified innate metabolic differences that arise with or in support of the resistant phenotype, despite large clinical variability, over multiple genetic backgrounds. Additionally, we were able to explore the functional consequences of expression changes by predicting essential enzymes within these distinct metabolic contexts; these enzymes are candidate drug targets for the prevention of drug resistance.

## RESULTS

### Analysis of Artemisinin Sensitive and Resistant Transcriptomes

In order to investigate the presence of a distinct metabolic phenotype in artemisinin resistant parasites, we analysed a previously published expression dataset of clinical isolates from Southeast Asia (NCBI Gene Expression Omnibus accession: GSE59097). Patient blood samples were collected immediately prior to beginning artemisinin combination therapy and relative expression was evaluated using microarrays [49]. We confined our analysis of this previously published expression data to ring-stage parasites from Cambodia and Vietnam, two countries that had clear resistant and sensitive parasite populations as defined by parasite clearance half-life, an *in vivo* phenotypic measure of resistance, and *Pf*Kelch13 mutations, a commonly-used genetic marker of resistance (Figure 1A & 1B). There were 97 and 24 ring-stage *resistant* parasite expression profiles from Cambodia and Vietnam, respectively; resistant parasites are defined by both the presence of *Pf*Kelch13 mutations and a parasite clearance half-life of more than 5 hours. There were 141 and 43 ring-stage *sensitive* parasite expression profiles from Cambodia and Vietnam, respectively, as defined by wild-type PfKelch13 alleles and clearance half-life of less than 5 hours. Despite obvious genotypic and phenotypic separation (Figure 1A & 1B), artemisinin sensitive and resistant parasites do not separate well by hierarchical clustering of expression data (Figure 1C).

**Figure 1:**
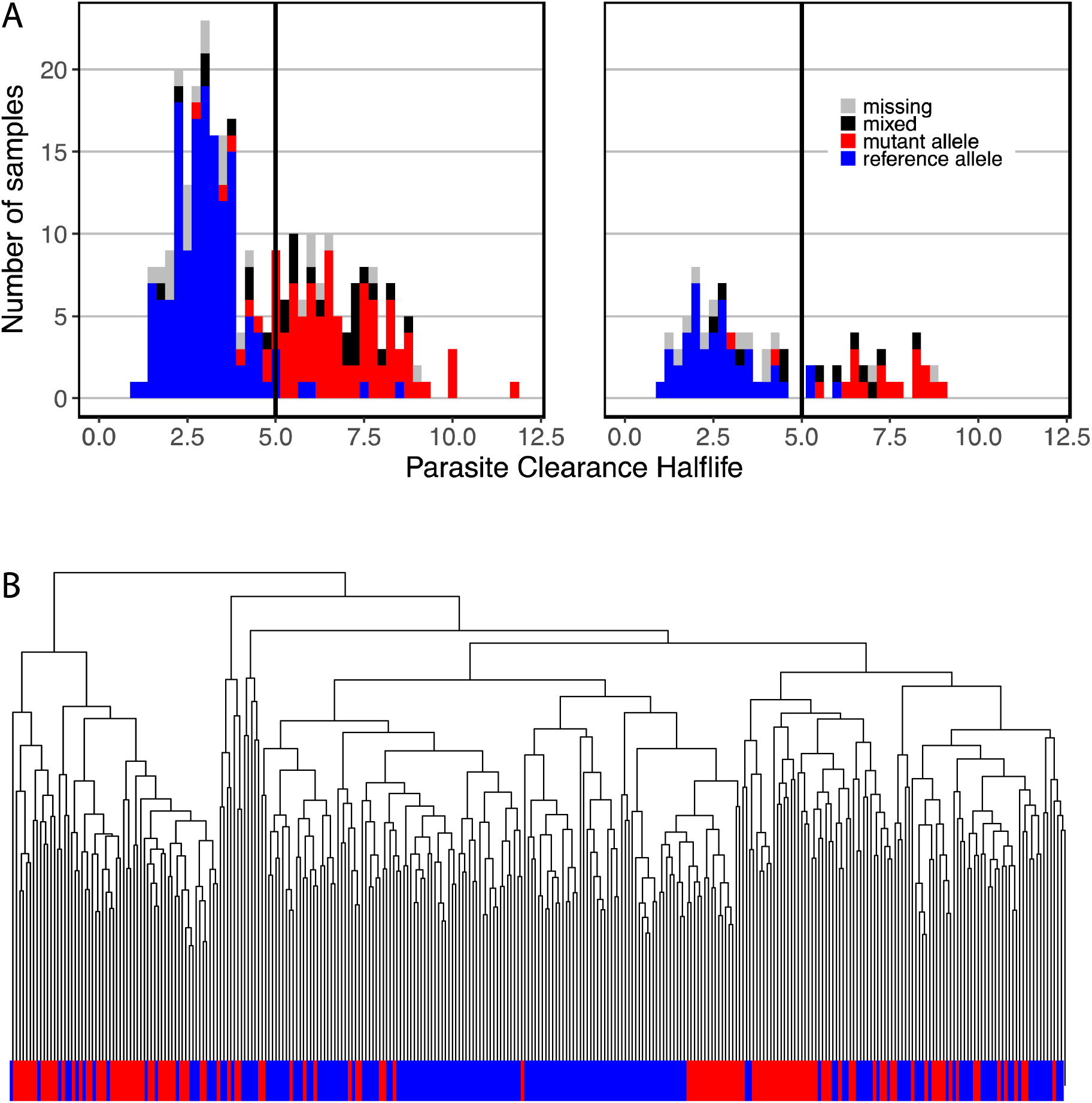
Ring-stage parasites are genotypically and phenotypically distinct, yet expression profiles fail to separate resistance phenotypes. ***A & B*** **Genotypic clustering**: Genotypic (any mutation in *Pf*Kelch13) and phenotypic markers (parasite clearance half-life) were used to define artemisinin resistance in ring-stage parasites from GSE59097; using both markers, resistant and sensitive parasites from Cambodia (**A**) and Vietnam (**B**) separated into distinct populations. ***C*** **Phenotypic clustering:** Resistant (*red*) and sensitive (*blue*) parasites fail to cluster with consideration of genome-wide gene expression data (data not shown) or expression of metabolic genes alone.

Additionally, fold change of the majority of transcripts is moderate; no genes exhibited notable differential expression across both analyses (fold change > 2 or < 0.5 for both Cambodia and Vietnam sample sets, data not shown). Among metabolic genes specifically, expression differences are small (maximum fold change 0.6 and 1.6) and few are both significant and conserved between data sets (11 in common from 174 in Cambodia and 37 in Vietnam; Suppl. Figure 1A & 1B). Large amounts of transcriptional variation (due to stage-dependent expression, genotypic variability, and host-pathogen interactions) across the population of clinical parasites may hide differences in the data sets. Moreover, we built a Random Forest classifier with expression data to predict resistance outcomes; the classifier predicted resistance poorly, with only 30.77% sensitivity (indicating only 30.77% of resistant samples were correctly identified) and 97.96% specificity (indicating 97.96% of sensitive samples were correctly identified) (Suppl. Figure 2A).

Although the expression data classifier performed poorly, a similar classifier built from metadata associated with each sample (patient and parasite characteristics) was highly predictive of resistance status with 85.71% sensitivity and 88.91% specificity (Suppl. Figure 2B). In our analysis, two specific mutations were the most predictive of resistance status, with sample collection site as the third most important variable; removing any of these three variables decreased classifier accuracy by over 20%. If Kelch13 mutations are used to predict resistance (rather than used to define resistance), Kelch13 mutations are most predicative of resistance (data not shown). Thus, metadata better predicts resistance than expression data. In order to deconvolve this innate variability and identify functional cellular changes associated with varying levels of artemisinin sensitivity, we integrated metabolic expression data into a genome-scale metabolic model of blood-stage *P. falciparum*.

### Manual metabolic network curation

To maximize the predictive ability of the metabolic network model, we curated an existing, well-validated reconstruction of asexual blood-stage *P. falciparum* [50] to improve its scope, and species-and stage-specificities. Our curated reconstruction, iPfal17, includes all metabolic reactions encoded by characterized genes in the parasite’s genome, summarizing metabolic behavior during the asexual blood-stage parasite. It is larger in scope from the previously published version due to the addition of 268 reactions (Table 1, **Suppl. Tables 1 & 2**), with 9.6% more enzymatic reactions and 2.3% more reactions with gene annotations. We also added 124 genes to the network (Table 1 & **Suppl. Table 1**). iPfal17 has gene annotations for 80.0% of enzymatic reactions, and 20.5% of transport and exchange reactions (Figure 2). iPfal17 includes 25.4% of the 1178 EC annotations in the *P. falciparum* genome, adding 14 EC numbers [43] (**Suppl. Table 1**). We evaluated enzyme complex or isozyme status and replaced 7 gene-protein-reaction relationships (**Suppl. Table 1**).

**Figure 2:**
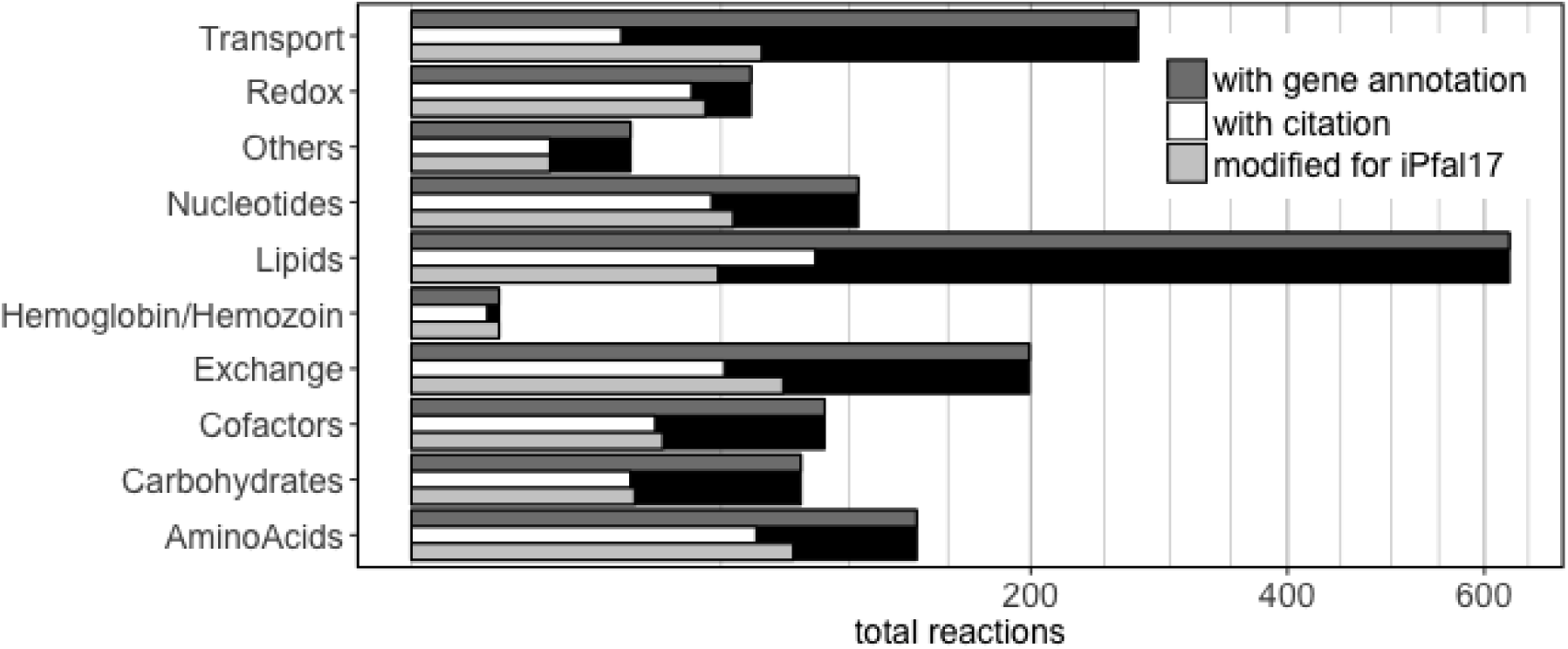
iPfal17 model curation is broad and comprehensive. Number of reactions in the *P. falciparum* reconstruction grouped by metabolic subsystems. Subsets of those reactions with gene annotations, literature citations, and modifications in the curation effort for this reconstruction are noted.

**Table 1:**
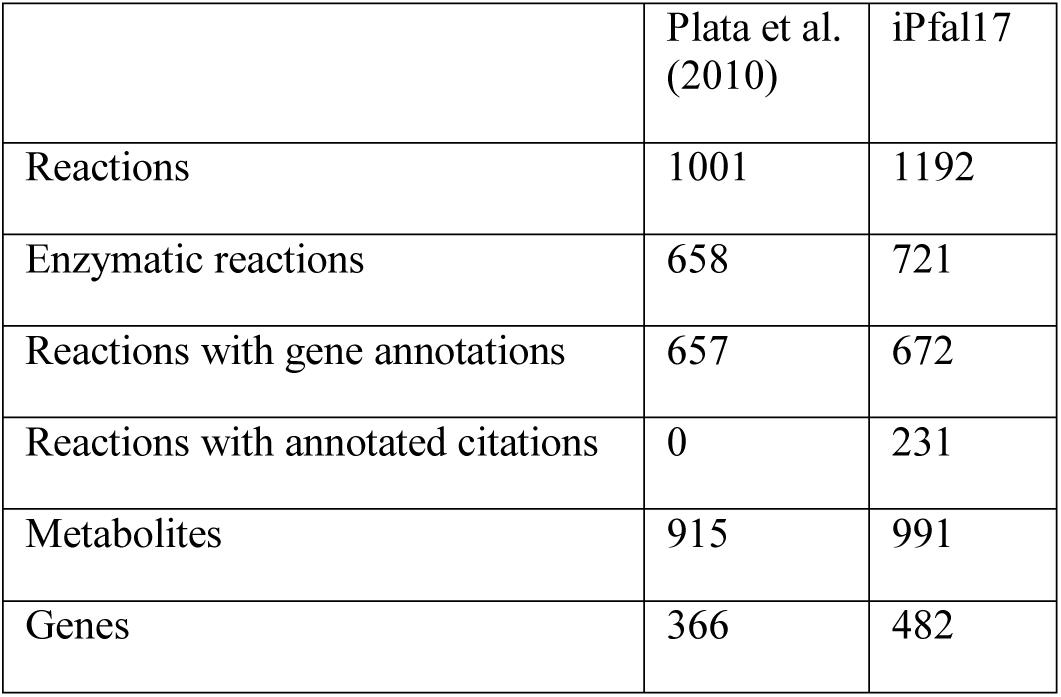
Asexual blood-stage *Plasmodium falciparum* parasite model, iPfal17, summary statistics.

Following curation, the species-and stage-specificity of the model was also improved. Gene annotations were evaluated against PlasmoDB resources [43], resulting in 124 additional gene annotations (**Suppl. Table 1)**. Importantly, we removed cellular import of pyrimidines from the host erythrocyte, as *P. falciparum* relies on *de novo* synthesis (**Suppl. Table 2)** [47, 51].

Blood-stage specificity was improved by removing genes only used in other life stages (specifically the gene encoding chitinase [52]). Additionally, 77 functionally unnecessary reactions were removed due to a lack of genetic and biochemical support (**Suppl. Table 2**). Reactions necessary for growth were added manually (**Suppl. Table 1**). Reactions were individually curated, changing metabolite utilization and stoichiometry (**Suppl. Table 1**).

The iPfal17 reconstruction contains five compartments: extracellular space and four intracellular compartments (cytoplasmic, mitochondrial, apicoplast, and food vacuole, **Suppl. Table 1)**. Few studies since the Plata et al. reconstruction investigated protein localization and therefore, few changes were made to compartmental assignments; the food vacuole compartment, containing two reactions, was added in this version of the reconstruction (**Suppl. Table 1**). As in the Plata et al. reconstruction, reactions with unknown localization were placed within the cytoplasm (**Suppl. Table 1**) [53]. Again, similar to Plata et al., a mitochondrial inner matrix was not added, as there is no evidence that the blood-stage parasite requires a proton gradient for energy production [51, 54, 55]. Nonpolar metabolites generated in one compartment and utilized in another were transported as needed for network functionality by assuming passive diffusion [53].

We also included annotations that will accelerate future curation efforts. First, we did not remove blocked reactions (those that do not carry flux due to their lack of connectivity to other components of the network) because further research may add connectivity to these network components. iPfal17 contains 303 blocked reactions and 78 dead-end metabolites (specifically, 32 metabolites are not consumed and 46 are not produced). For example, 4-pyridoxate (a byproduct of vitamin B6 biosynthesis) is included; production is supported by bioinformatic analyses of the parasite genome, but the metabolite function or excretion pathway is not known. Second, citations are included within iPfal17 to identify the date of discovery and degree of literature support for each reaction (**Suppl. Table 1 & 2**). Literature support was only added to modified reactions, resulting in 231 citations (Table 1 & **Suppl. Table 1**).

### Metabolomics curation of biomass reaction

For the newly curated iPfal17 model, we modified the *Plasmodium* biomass reaction to better represent *in vitro* data (Table 2). We added tRNA-ligated amino acids to the amino acid requirements to force protein production, rather than only demanding free amino acids. Additionally, lipid classes were added based on recently published metabolomics findings; phosphatidylinositol, phosphatidyl-glycerol, sphingomyelin, diacylglycerides, and triglycerides were added due to their observed increase in abundance between uninfected and infected erythrocytes [56]. Phosphatidylcholine ethers, acyl phosphatidylgycerol, lyso-phosphatidylinositol, bis(monoacyl-glyceryl)phosphate, and monosialodihexosylganglioside were excluded from the biomass reaction, as there is no known *Plasmodium* catabolism or import pathways for these lipids [56]. Analysis of metabolomics data enabled further curation of the biomass reaction with the addition of malate, alpha-ketoglutarate, and glutathione (both reduced and oxidized) [28, 57, 58]. Importantly, we included the requirement for cellular export of lactate and hemozoin. Lactate is measured in extracellular *in vitro* metabolomics and *in vivo* via blood acidosis; it is the terminal product of glycolysis, the sole energy production pathway used by the blood-stage parasite [59-62]. By requiring lactate export, we force the model to utilize glycolytic energy metabolism. Similarly, hemoglobin degradation is essential for the blood-stage parasite to produce free amino acids. Parasites can also import and synthesize some amino acids, but the breakdown of hemoglobin (and subsequent production of its byproduct, hemozoin) is necessary for growth [23, 63, 64]. Thus, by requiring hemozoin export, we force the *in silico* parasite to degrade hemoglobin as the primary pathway for amino acid production.

**Table 2:** Metabolic components of the biomass function.

|  |  |  |
| --- | --- | --- |
| **Complex metabolites** | • protein<br>• lipid<br>• reduced and oxidized glutathione<br>• protoheme |  |
| **Amino Acids** | • alanine<br>• asparagine<br>• cysteine<br>• glutamine<br>• histidine<br>• leucine<br>• methionine<br>• phenylalanine<br>• threonine<br>• tyrosine | • arginine<br>• aspartate<br>• glutamate<br>• glycine<br>• isoleucine<br>• lysine<br>• serine<br>• proline<br>• tryptophan<br>• valine |
| **Carbohydrates** | • malate<br>• α-ketoglutarate |  |
| **Nucleotides** | • ATP<br>• CTP | • dATP<br>• dCTP |
|  | • GTP<br>• UTP<br>• thiamine diphosphate | • dGTP<br>• dTTP |
| **Excreted metabolites** | • lactate<br>• hemozoin |  |
| **Vitamins** | • pyridoxal<br>• 5-phosphate riboflavin |  |
| **Other** | • s-adenosyl l-methionine<br>• spermidine<br>• f-thf<br>• thf<br>• coenzyme-A<br>• water<br>• NADP<br>• NH<sup>4+</sup> | • putrescine<br>• 2-octaprenyl 6-hydroxyphenol<br>• mthf<br>• FAD<br>• NAD<br>• Fe<sup>2+</sup> & Fe<sup>3+</sup><br>• SO<sup>4</sup> |

f-thf = formyl tetrahydrofolate; mthf = methyltetrahydrofolate; thf = tetrahydrofolate

### iPfal17 validation and functional requirements

To validate the model against experimental results, essential metabolic tasks of blood-stage growth were identified and evaluated (Table 3). These tasks describe experimental and clinical observations, such as the parasite’s ability to grow on glucose and hypoxanthine as a sole carbon source and purine source, respectively, and the parasite’s induction of blood acidosis via lactate [65-67]. Additional tasks include the parasite’s failure to grow in the presence of antimetabolites for riboflavin, nicotinamide, thiamine, and pyridoxine [68]. We defined this set of tasks to provide a framework for curation and validation efforts of future network reconstructions. Although iPfal17 fails to pass all metabolic tasks, we believe this is the most comprehensive and accurate model to date due to the curation efforts and results from tests of the metabolic tasks. Failures generally exist in pathways that currently contain many reversible reactions (i.e. tasks 5a-b for glycolysis) or if the experimental evidence is not mechanistic (i.e. tasks 1a-d) or fully characterized (i.e. task 4; Table 3).

**Table 3:** Experimentally derived metabolic tasks for evaluating iPfal17.

|  | Metabolic Task | *In vitro* | iPfal17 | Hypothesis for *in vitro/in silico* discrepancies |
| --- | --- | --- | --- | --- |
| 1a | Growth in the presence of antimetabolite, riboflavin? | no [68] | no | - |
| 1b | Growth in the presence of antimetabolite, thiamine? | no [68] | yes | Unknown antimetabolite mechanism; Off target effects of antimetabolite |
| 1c | Growth in the presence of antimetabolite, nicotinamide? | no [68] | yes | Unknown antimetabolite mechanism; Off target effects of antimetabolite |
| 1d | Growth in the presence of antimetabolite, pyridoxine? | no [68] | yes | Unknown antimetabolite mechanism; Off target effects of antimetabolite |
| 2a | Grows without loops? | no | no | - |
| 2b | ATP production if no exchange is allowed? | no | no | - |
| 3a | Can produce purines? | yes | yes | - |
| 3b | Growth with hypoxanthine as the only purine source? | yes [65] | yes | - |
| 3c | No growth if guanine, guanosine, inosine, adenine, or adenosine are only purine sources? | yes [65] | 60% | - |
| 4 | Growth with IPP supplementation and no apicoplast? | yes [69] | no | Nuclear encoded proteins that function within the apicoplast may function in the cytoplasm if the organelle is not present. |
| 5a | Growth with glucose? | yes [66] | yes | - |
| 5b | Growth with alternative sugar source (no glucose, with ribose, mannose, fructose, galactose, or maltose)? | no [66] | yes | Central carbon metabolism contains many reversible reactions. |
| 6a | Can produce all amino acids except isoleucine? | yes [66] | yes | - |
| 6b | Is growth reduced without methionine, proline, tyrosine, cystine, glutamate, or glutamine supplementation? | yes [63] | no | Model is not designed for growth reduction experiments. |
| 6c | Growth without isoleucine supplementation? | no [63, 70] | no | - |
| 7 | Growth without calcium pantothenate? | no [70] | no | - |
| 8 | Growth without p-aminobenzoic acid? | no [71] | no | - |
| 9 | Cannot produce any metabolites if no exchange is allowed? | no | no | - |
| 10 | Accuracy of experimental essentiality predictions | - | 79.5% | See Table 4 and Suppl. Table 7 |
| 11 | Accuracy of *P. berghei* essentiality predictions | - | 61.4% | See Suppl. Table 6 |

\* Accuracy calculated as the sum of true positives and true negatives, divided by total observations

We also evaluated predictions of the effects of gene knockouts and enzyme inhibitors using previously published experimental results (Table 4; **Suppl. Table 7**). Our updated model had improved accuracy of gene and reaction essentiality predictions, compared to previous models and specifically Plata et al. (Table 4). We predict that there are 159 essential reactions, and 107 lethal single gene knockouts (**Suppl. Table 3 & 4**). Of experimentally validated knockouts, iPfal17 accurately predicts essentiality of 79.5% of genes and enzymes tested in *P. falciparum* and 61.4% for those tested in *P. berghei* (Tables 3 & 4, **Suppl. Table 6**); predictions are also more accurate for gene knockouts and are less accurate in predicting enzyme inhibition (Tables 3 & 4).

**Table 4:**
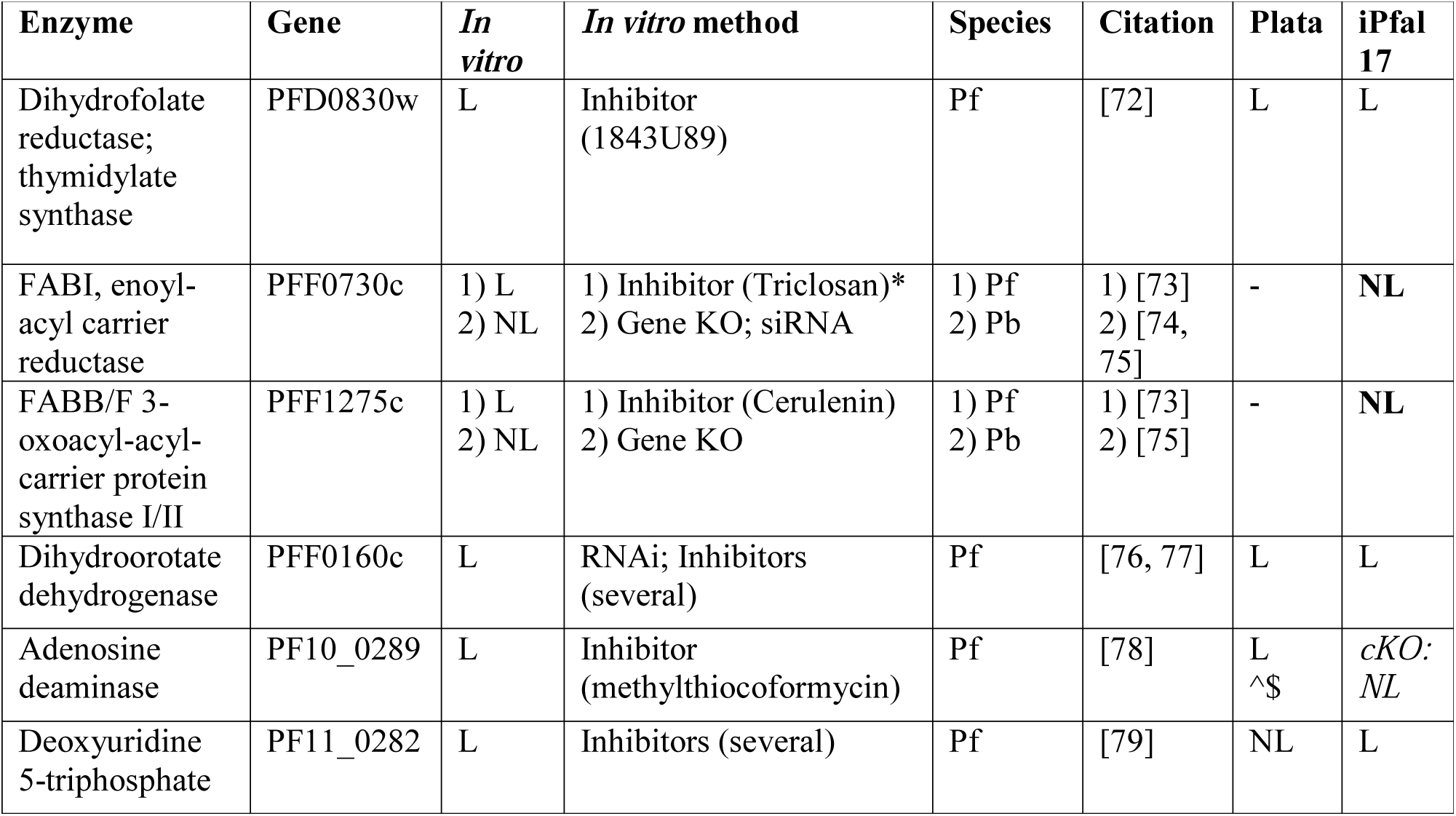
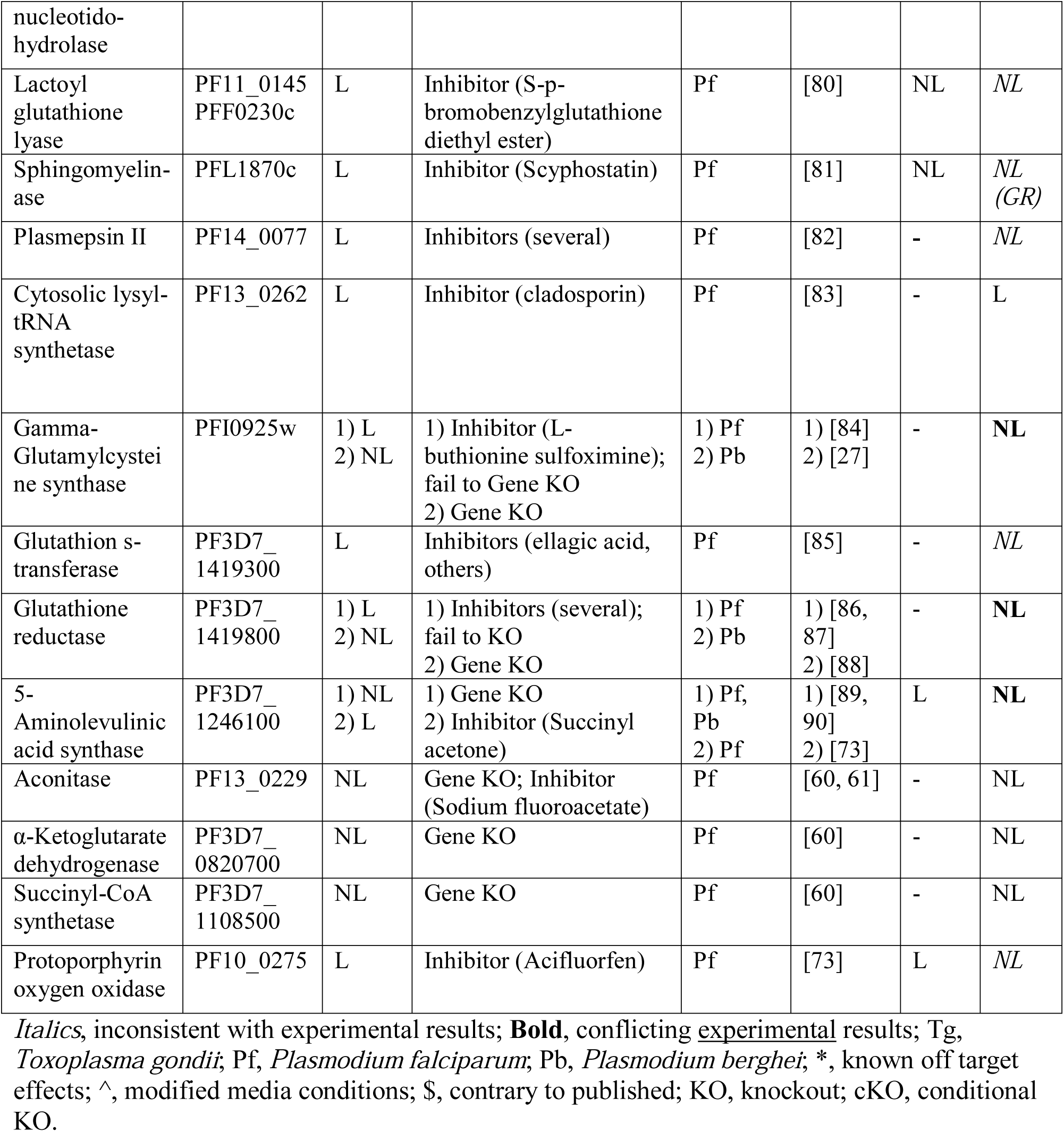
Knockout predictions with experimental validation. Predictions for 18 enzymes of interest are included here. See Suppl. Table 7 for the complete list of predictions.

### Integration of expression data into the metabolic model

With this high-quality metabolic network reconstruction, we integrated expression data from sensitive and resistant parasites collected in Cambodia and Vietnam into iPfal17 using the Metabolic Adjustment for Differential Expression algorithm (MADE [91]). MADE constrains gene utilization in the model to maximally account for statistically significant changes in expression data while accounting for network functionality requirements.

MADE integration of sensitive and resistant expression data from both countries generated four condition-specific models (Figure 3). Genes (and associated reactions) are removed from condition-specific models if their transcripts are down-regulated and not functionally necessary for metabolism in that condition. Those remaining in the constrained model are a subset of genes annotated in our original curated reconstruction; these genes are either more highly expressed in the corresponding condition, not differentially expressed across conditions, or necessary for network functionality, and, thus, remain in the condition-specific model. By comparing these models, we identified differences in gene and pathway utilization between resistant and sensitive parasites that are consistent between the isolates from the two countries (Suppl. Figure 3).

**Figure 3:**
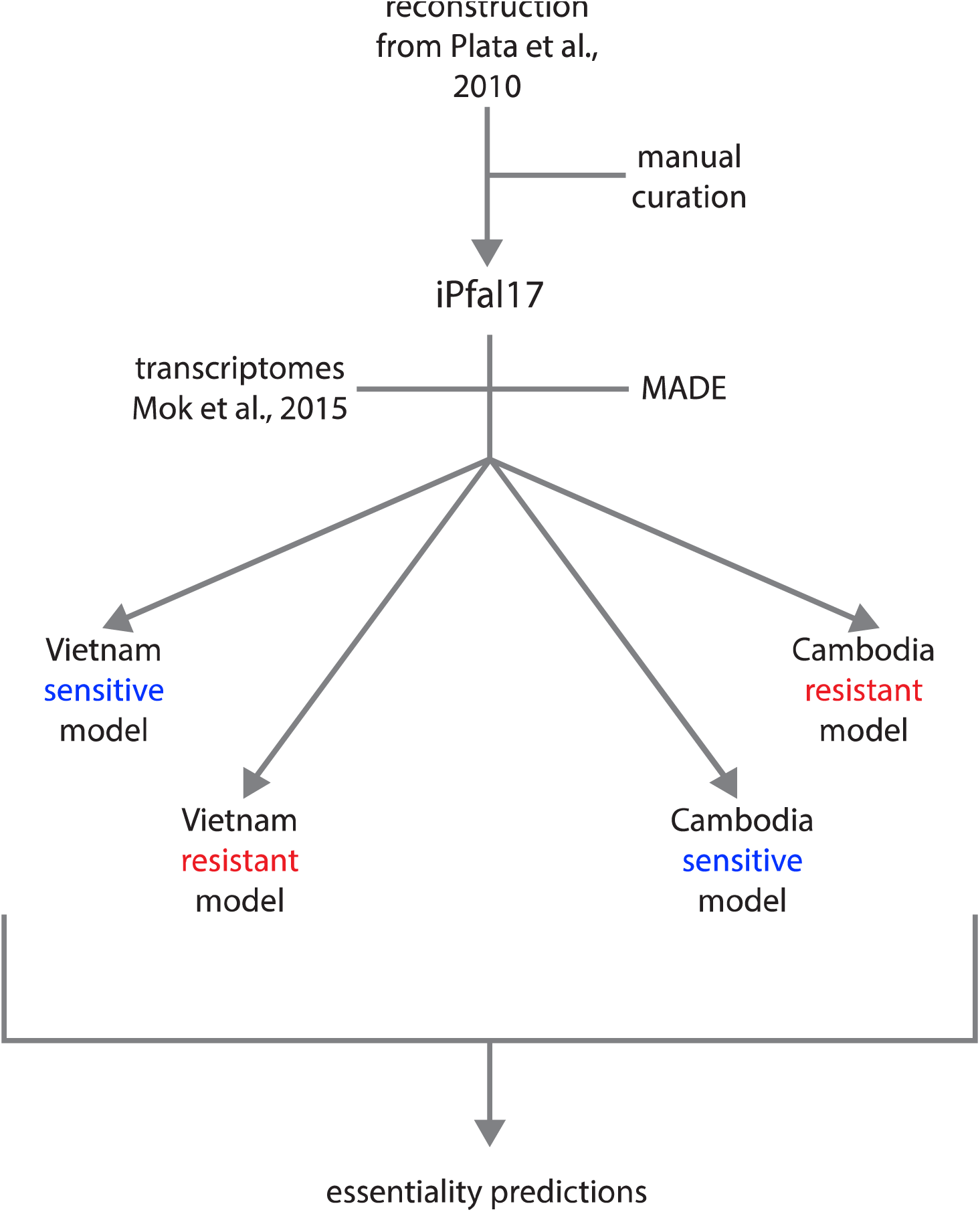
Computational pipeline. We curated an existing blood-stage *P. falciparum* reconstruction to generate our iPfal17 network reconstruction. We integrated transcriptomics data into this model using the MADE algorithm to generate four condition-specific models. We used these models to predict reaction essentiality; we highlight consensus results across resistant or sensitive models. MADE, Metabolic Adjustment for Differential Expression.

First, we conducted an enrichment analysis on genes that remain in (i.e. can be utilized by) each constrained model by comparing to the unconstrained curated model. As expected, all four models were enriched with genes involved in pathways with many essential reactions or little redundancy, such as transport reactions, tRNA synthesis, purine metabolism, and others (Suppl. Figure 4, see model). Sensitive, wild type, models corresponding to isolates from both Cambodia and Vietnam are uniquely enriched with the utilization of genes involved in the metabolism of nicotinate/nicotinamide (p-value = 1.47*10^-2^), glutamate (p-value = 1.28*10^-13^), and selenocysteine (p-value = 5.85* 10^-4^). Thus, sensitive models contain more reactions in these pathways than the unconstrained model, resulting from increased expression of these pathways in sensitive parasites (Suppl. Figure 4). Resistant models from both countries are uniquely enriched with the utilization of genes involved in pyrimidine (p-value = 2.18*10^-7^), polyamine (p-value = 4.39*10^-4^), redox reactions (p-value = 5.13*10^-5^), and central carbon metabolism (glycolysis [p-value = 4.39*10^-4^] and the pentose phosphate pathway [p-value = 6.06*10^-3^]). Thus, resistant models have a larger proportion of their total reactions associated with these pathways than the original unconstrained model, whereas sensitive models do not have this enrichment. This indicates that these pathways are upregulated in resistant parasites and may remain important for metabolism in the resistant state (Suppl. Figure 4).

**Figure 4:**
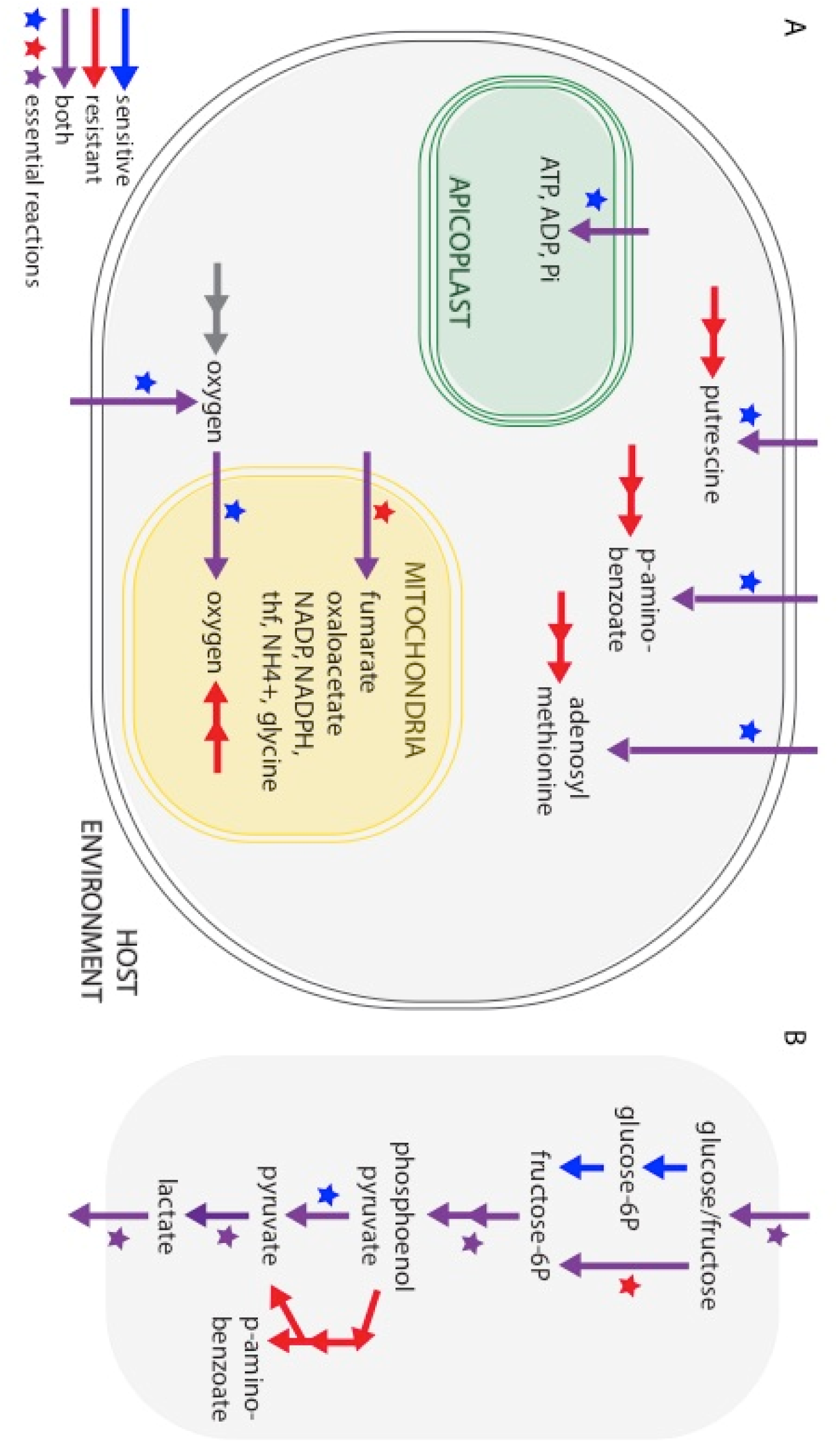
Artemisinin resistant and sensitive parasites have unique metabolite transport capabilities. **A Transport differences**: Resistant parasites exhibit greater metabolic flexibility, allowing either import or biosynthesis of putrescine, p-aminobenzoate, adenosyl-methionine into the parasite’s cytoplasm (*grey*). Sensitive parasites rely on import only Import or synthesis of ATP, ADP, and phosphate into the apicoplast (*green* organelle) is essential for sensitive parasites. Resistant parasites require transport of oxygen, fumarate, oxaloacetate, NADP, NADPH, tetrahydrofolate (thf), NH_4_, and glycine into the mitochondria, in *yellow. B* **p-aminobenzoate in glycolysis**: Resistant parasites generate p-aminobenzoate via alternative components of the glycolysis pathway. Arrows colored for flux via FVA and stars for essentiality. FVA, flux variability analysis.

### Identification of conserved and uniquely essential pathways

Beyond general differences in pathway utilization, which encompasses both essentiality and pathway-level differences in expression, artemisinin sensitive and resistant parasites have unique essential genes and reactions. To identify these essential reactions and provide insight on targetable metabolic enzymes in the clinical isolates, we performed *in silico* single gene and reaction deletions with each of the four condition-specific models. Datasets from the parasites from each country were initially analyzed separately and then lists were compared to ensure resistance-associated trends are reproducible and observed in independent analyses. As expected, we identified many essential functions conserved in all models (**Suppl. Table 5**), which is consistent with an active core metabolism required for basic parasite survival. Importantly, 21 reactions were essential in only resistant models, but not sensitive models (Table 5). Theoretically, drugs targeting these reactions would kill resistant parasites and have no effect on sensitive parasites; thus, there would be no selective pressure within the sensitive parasite population to develop resistance to these drugs. This list included serine hydroxymethyltransferase (PFL1720w in folate metabolism), the glycine cleavage system (PFL1550w and others in folate metabolism), thiamine diphosphokinase (PFI1195c in cofactor metabolism, specifically thiamine diphosphate), fumarate hydratase and malate dehydrogenase (PFI1340W and PFF0895w, respectively, in the mitochondrial electron transport chain and TCA cycle), and fructose hexokinase (PFF1155W in glycolysis; Table 5). We also identified 12 reactions that were essential only in artemisinin sensitive parasites (Table 6). Drugs targeting these reactions should not be combined with artemisinin, as they would not kill (and may select for) resistant parasites. Fortunately, no existing drug targets were found in this list of essential genes and reactions (Table 6). Among those identified were sphingomyelin synthase 2 (PFF1215w) and several transport reactions, which furthers our understanding of condition-specific intra-organellar function (Table 5 & 6). Overall, our systems biology-based approach reveals unique metabolic phenotypes associated with artemisinin sensitivity; these differences were not detected in the original analysis of the expression dataset or by separately analyzing Cambodian or Vietnamese isolates ([49] and data not shown).

**Table 5:** Essential reactions unique to resistant parasites. All reactions in table are predicted to be lethal when removed from both Cambodia and Vietnam resistant models.

| Reaction | Reaction Formula | EC Number | Reaction Function | Genes |
| --- | --- | --- | --- | --- |
| CO2tmt | CO<sub>2</sub>[m] <=> CO<sub>2</sub>[c] | -- | CO<sub>2</sub> transport | - |
| EX_folate4 | p-aminobenzoate[e] <=> | - | p-aminobenzoate exchange | - |
| EX_fru(e) | fructose[e] <=> | - | fructose exchange | - |
| EX_thm(e) | thiamine[e] <=> | - | thiamine exchange | - |
| FRUt1r | fructose[e] <=> fructose[c] | - | fructose transport | PFB0210c |
| FUM_mt | fumarate [m] + H<sub>2</sub>O[m] <=> malate[m] | 4.2.1.2 | fumarate hydratase in the TCA cycle | PFI1340w |
| FUMtmt | fumarate[m] <=> fumarate [c] | - | fumarate transport into mitochondria | - |
| GHMT2r | serine[c] + thf[c] <=> glycine[c] + H<sub>2</sub>O[c] + mthf[c] | 2.1.2.1 | serine hydroxymethyltransferase in folate synthesis | PFL1720w |
| GLYCL_mt | glycine[m] + NAD[m] + thf[m] <=> CO<sub>2</sub>[m] + mlthf[m] + NADH[m] + NH<sub>4</sub>[m] | many | glycine cleavage system in folate synthesis and amino acid metabolism | PF13_0345*<br>PFL1550w<br>MAL13P1.390<br>PF14_0497<br>PF11_0339 |
| GLYtmt | glycine[m] <=> glycine[c] | - | glycine transport into mitochondria | - |
| HEX7 | ATP[c] + fructose[c] => ADP[c] + fructose-6-phosphate[c] + h[c] | 2.7.1.1 | hexokinase of glycolysis | PFF1155w |
| MDHm | malate[m] + NAD[m] <=> h[m] + NADH[m] + oxaloacetate[m] | 1.1.1.37 | malate dehydrogenase in the TCA Cycle | PFF0895w |
| MLTHFtmt | mthf[m] <=> mthf[c] | - | mthf transport into mitochondria | MAL8P1_13*<br>PF11_0172 |
| NADPHtmt | NADPH[c] <=> NADPH[m] | - | NADPH transport into mitochondria | - |
| NADPtmt | NADP[c] <=> NADP[m] | - | NADP transport into mitochondria | - |
| NH4tmt | NH<sub>4</sub>[m] <=> NH<sub>4</sub>[c] | - | NH<sub>4</sub> transport into mitochondria | - |
| OAAtmt | oxaloacetate [m] <=> oxaloacetate[c] | - | oxaloacetate into mitochondria | - |
| THFtmt | thf[m] <=> thf[c] | - | thf into mitochondria | - |
| THMt3 | h[c] + thiamine[e] <=> h[e] + thiamine[c] | - | thiamine import | - |
| TMDPK | ATP[c] + thiamine[c] => AMP[c] + h[c] + thiamine diphosphate[c] | 2.7.6.2 | thiamine diphosphokinase in cofactor metabolism | PFI1195c |
| pABAt | p-aminobenzoate[e] <=> p-aminobenzoate [c] | - | p-aminobenzoate import | MAL8P1_13*<br>PF11_0172 |

\*, deleted from at least one resistant model due to expression constraints by MADE localization: [e] extracellular, [c] cytoplasmic, [m] mitochondria, [ap] apicoplast

**Table 6:**
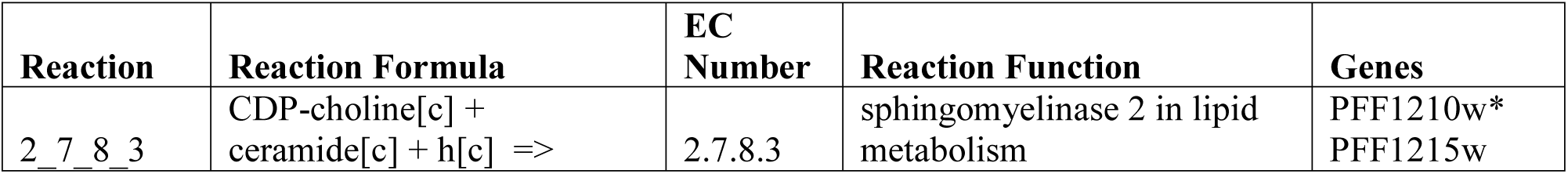
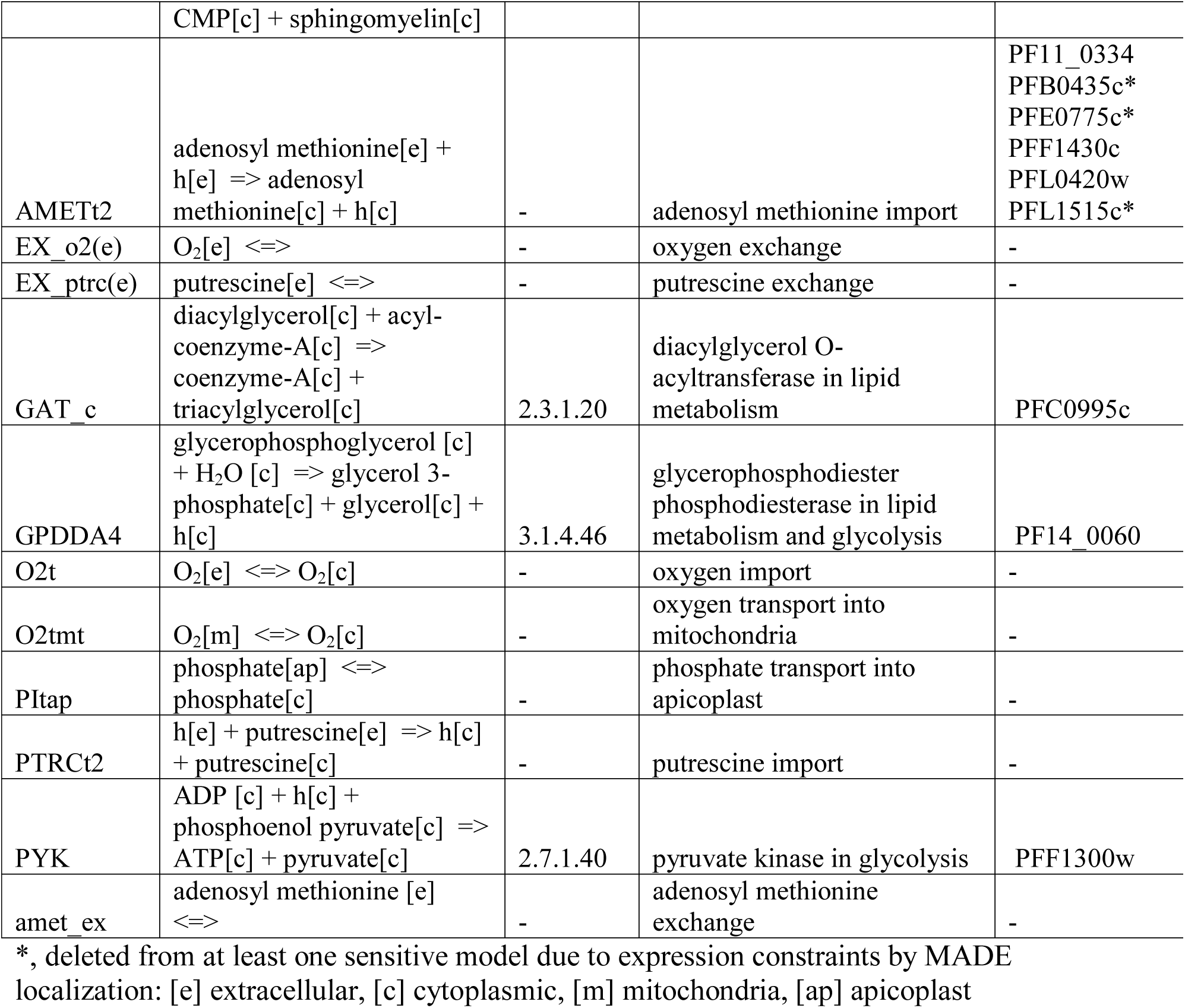
Essential reactions unique to sensitive parasites. All reactions in table are predicted to be lethal when removed from both Cambodia and Vietnam sensitive models.

## DISCUSSION

Systems biology approaches enable unbiased analyses of antimalarial resistance phenotypes. Here, we describe a newly curated metabolic network reconstruction of the malaria parasite that can serve as a platform for the analysis of gene expression and other ‘omics data, and as a tool to generate testable hypotheses regarding essential genes and metabolic phenotypes. In particular, we used this network reconstruction to characterize key metabolic dependencies in resistant and sensitive parasites. We revealed emergent patterns in pathway activity, differential utilization of organelles, metabolic flexibility, and targetable weakness of resistant parasites.

### Data-driven model curation improves predictive capability

iPfal17 represents the most comprehensive, stage-specific *P. falciparum* metabolic reconstruction to date. With iPfal17, we can simulate growth and predict gene and reaction essentiality. It is larger in scope than previous models, includes more gene annotations, and documents literature citations associated with its components (Table 1 & **Suppl. Table 1**, Figure 2). Moreover, invalid reactions have been removed, improving accuracy (**Suppl. Table 2**). These curation efforts improve the model validity by better recapitulating experimental results, removing functions known to not occur in the asexual blood-stage parasite, and adding functions for which there is experimental evidence. Thus, gene and reaction knockout predictions generated with this model are more accurate. Moreover, iPfal17 has greater interpretability as reaction citations and confidence scores are included and accessible to users.

iPfal17 is similar in functional distribution and scope to other high quality models of apicomplexans, despite its reduced genome size. The *P. falciparum* genome is 23.3MB and contains 5423 genes (excluding the antigenic var genes) [92] [43, 93]; iPfal17 accounts for the function of 987 metabolites, 730 enzymatic reactions, 1195 total reactions, and 488 genes (**Table 1**). For reference, the network reconstruction for *Toxoplasma gondii*, with a genome of 80 MB with 8,000 genes [94, 95], accounts for 1019 metabolites, 1089 enzymatic reactions, 3387 total reactions, and 527 genes [96, 97]; with a genome of 32.8 MB and 8272 genes [97], the *Leishmania major* network reconstruction accounts for 1101 metabolites, 1047 enzymatic reactions, 1112 total reactions, and 560 genes [98]. These parasites all have notably poor genome annotation (40-60% of the genes are unknown) [43, 93] and, thus, have fewer associated genes than many other reconstructions (e.g. the *E. coli* and *S. cerevisiae* reconstructions account for 1366 and 910 genes[99], respectively [99, 100]).

Intracellular parasites, like *Plasmodium*, require more exchange and transport reactions as they obtain many nutrients from the host environment [57, 101-104]. This reliance on the host for metabolic function permits the parasite to increase fitness by reducing its genome and hijacking host function. *P. falciparum* does just that: the parasite remodels the host erythrocyte, generating a vesicular network for protein translocation and increasing host cell permeability for nutrient acquisition from the host serum [105-109]. Thus, the apicomplexan network reconstructions include more transport reactions, many of which are not genetically mapped. Additionally, we chose to exclude an erythrocytic host compartment from the extracellular environment, despite the parasite’s intra-host growth [57, 66, 68, 70]. Other recent reconstructions [110, 111] have added this compartment, but the erythrocytic compartment is unlikely to improve model function due to the gross disruption of the host membrane as a barrier [57, 66, 68, 70].

We generated gene and reaction essentiality predictions with our curated network model, prior to integration of expression data, and found results largely consistent with previous models [50] (Table 4). We identified 159 essential reactions and 107 essential metabolic genes (**Suppl. Table 3 & 4)**; 24 of these have been empirically tested in cultured *P. falciparum* parasites (Table 4, and in *P. berghei-* **Suppl. Table 6**). iPfal17 better predicts experimentally determined essential reactions than previous models, across a broad set of metabolic pathways (Table 4 and data not shown). iPfal17 predictions fail when essential genes or reactions are involved closely with spontaneous reactions (i.e. lactoylglutathione lyase is downstream of a spontaneous reaction and upstream of nonmetabolic redox products), are in pathways with uncharacterized mechanisms (i.e. plasmepsin II in hemoglobin degradation) or if experimental evidence is contradictory (i.e. heme biosynthesis pathway; **Table 4**).

Because pharmacological enzyme inhibition can be quite noisy and genetic modification has been challenging in *Plasmodium*, the development of CRISPR-Cas9 and other technologies will make it possible to integrate new experimental observations into the model with increasing accuracy [112-115]. Until then, the model can be used to identify enzyme inhibitors with off-target effects. For example, within the heme biosynthesis pathway, pharmacological inhibition of aminolevulinic acid dehydrogenase and protoporphyrinogen oxidase kills blood-stage parasites [73]; however, disrupting the genes encoding the first (aminolevulinic acid dehydrogenase) and last (ferrochetalase) genes is not lethal in blood-stage parasites [89, 90]. iPfal17 predictions are consistent with the gene knockout experiments in *P. falciparum* of Ke, et al., suggesting that the enzyme inhibitors used by Ramya, et al. have off target effects (**Table 4** and **Suppl. Table 7**). iPfal17 also fails to predict the lethal nature of adenosine deaminase in purine-free conditions [78]. Adenosine deaminase converts adenosine to hypoxanthine; as 38 reactions produce AMP, which then generate hypoxanthine products, we propose adenosine deaminase may be essential for nonmetabolic functions or the inhibitor of adenosine deaminase has off target effects. Furthermore, these results generate hypotheses about the differential metabolic capabilities of *P. falciparum* and *P. berghei*, as experimental results in the rodent parasite conflict with some *P. falciparum* predictions (**Tables 3 & 4, Suppl. Table 6**).

### Data integration reveals distinct metabolic patterns

The integration of expression data from clinical parasites into our network reconstruction highlights the differential utilization of metabolic genes and reveals metabolic shifts associated with variation in innate artemisinin sensitivity (Suppl. Figure 3 & 4). Enriched metabolic pathways detected in sensitive and resistant models are consistent with previous experimental observations. For example, resistant models are uniquely enriched with genes involved in pyrimidine biosynthesis and mitochondrial redox reactions. This finding is consistent with the importance of mitochondrial function in surviving artemisinin stress [26, 29]. Additionally, the metabolic disruption of the redox reactions in the electron transport chain upon artemisinin treatment (via decreased production of orotate and fumarate, presumably via dihydroorotate dehydrogenase and succinate dehydrogenase enzymes [22, 28, 116]) suggests that changes in these pathways may be important for survival in the presence of the drug. Thus, this metabolic network analysis approach allows us to filter out noise from diverse clinical isolates to identify alternative utilization of pathways associated with artemisinin resistance. However, these enrichment results do not implicate specific reactions that are uniquely active in artemisinin sensitive or resistant parasites.

### Condition-specific models have unique metabolic requirements

Upon integration of expression data and the identification of differentially utilized pathways above, we next used these models to predict targetable differences in sensitive and resistant parasites by identifying reactions that are essential within the context of the metabolic network (**Suppl. Tables 5, Tables 5 & 6**). We identified (1) differences in intra-organellar function, (2) metabolic flexibility of scavenging and biosynthesis pathways, and (3) targetable weakness of resistant parasites. These metabolic shifts primarily reside in mitochondrial metabolism, as well as folate and polyamine metabolism. Together, these results highlight the overall plasticity of *P. falciparum* metabolism and opportunities for further development of potential drug targets.

Interestingly, several transport reactions are found to be differentially essential in our constrained models (Tables 5 & 6). Many transport reactions (79.5%) have no associated gene due to the incomplete characterization of the *P. falciparum* genome (Figure 2). They are included in the model due to biochemical evidence or functional necessity (i.e. a metabolite is produced in one compartment but it is a substrate for an enzyme in another). Transcriptomic data integration does not constrain their behavior explicitly: expression integration reduces the total number of reactions in a model, forcing transport of metabolites among organelles if within-compartment biosynthesis is non-functional. Function within organelles requires transport and loss of function reduces transport needs. Specifically, several mitochondrial and apicoplast transport reactions are uniquely essential in the sensitive and resistant parasite populations (Figure 4). In resistant models, this includes the mitochondrial transport of metabolites associated with the TCA cycle and electron transport chain (fumarate, oxaloacetate, and NADPH) and those involved in generation of folates (tetrahydrofolate, glycine, CO_2_, and NH_4_+) (Figure 4A). In sensitive models, apicoplast transport of ADP, ATP, and phosphate is essential (Figure 4A). Overall, these results indicate that sensitive and resistant parasites are differentially utilizing pathways within these organelles, and have unique requirements for transport of essential substrates. This observation is consistent with previous studies and our enrichment results highlighting the influence of mitochondrial metabolism on survival in the presence of artemisinin [26, 29]. Moreover, oxygen transport into the cell and then into the mitochondria is only essential in sensitive parasites, further predicting differential use of the mitochondria in these parasites as oxygen serves as the terminal step in the electron transport chain. Resistant parasites are predicted to generate oxygen within the mitochondria via superoxide dismutase as opposed to transport (Figure 4A).

We also identify differential utilization of transport pathways from the extracellular environment into the parasite. *Plasmodium* metabolism contains redundancies; for many essential metabolites, the parasite’s genome encodes one or more biosynthetic pathways, while there is also evidence for a parallel host-scavenging pathway [108] (e.g. lipid [56] and amino acid [56, 63] scavenging). Upon model integration, we find that artemisinin resistant and sensitive parasites utilize some of these metabolic pathways in alternative ways (Figure 4A). *Plasmodium* can either scavenge or synthesize putrescine and adenosyl-methionine (essential polyamines and precursors to spermidine [50, 117]), as well as p-aminobenzoate, a folate precursor generated by branch of glycolysis necessary for nucleotide synthesis ([118]; Figure 4 A&B, Tables 5 & 6). These metabolites are measurable via blood sample metabolomics [118, 119]; therefore, host scavenging is a viable option for blood-stage parasites. We predict that sensitive parasites rely on the import of putrescine, adenosyl methionine, and p-aminobenzoate. Resistant parasite expression supports either this host scavenging or direct biosynthesis due to parasite survival upon reaction knockout *in silico*. Thus, we expect that resistant parasites are more metabolically flexible for these metabolites; perhaps resistant parasites have failed to appropriately modulate their transition to the nutrient-rich blood-stage environment, and this unexpected flexibility is evolutionarily beneficial once confronted with artemisinin treatment.

Interestingly, recent metabolomics studies demonstrate that intra-parasitic putrescine levels are decreased drastically upon artemisinin treatment [116]; activity in both the biosynthetic and scavenging pathway of putrescine and adenosyl methionine may allow resistant parasites to compensate for artemisinin’s effect on polyamines. The essential role of polyamines is well established in *Plasmodium* [120, 121]. In other organisms, these compounds stabilize DNA and RNA [122] and signal a pause in the cell cycle [123]. In the presence of artemisinin, perhaps polyamines act to stabilize the genome from oxidative stress [24, 30, 33] and trigger dormancy [18, 19]. As resistant parasites are more likely to survive dormancy, flexibility in polyamine metabolism could provide more routes for artemisinin survival [29, 124].

Our systems biology approach also identifies metabolic weaknesses of resistant parasites; these weaknesses can be used to identify drug targets for combination therapies (Figure 5). For example, we identified the mitochondrial import of fumarate and subsequent conversion to oxaloacetate (via fumarate hydratase, PFI1340W, and malate dehydrogenase, PFF0895W) to be uniquely essential in resistant parasites (Figure 5A, Table 5). Expression data from sensitive parasites supports mitochondrial import of malate and utilization of malate:quinone oxidoreductase (PFF0815W) to generate oxaloacetate from malate, bypassing the need for fumarate and the associated enzymes, fumarate hydratase and malate dehydrogenase. We predict that inhibitors of fumarate transport or fumarate hydratase and malate dehydrogenase would specifically kill artemisinin resistant parasites, offering an example of enhanced metabolic flexibility of sensitive parasites and a potential artemisinin-combination therapy target. The TCA cycle is essential during the mosquito-stage of parasite development [61, 125], but not the blood-stage [60, 61]; this once again highlights the possibility that resistant parasites exhibit incomplete transition to the metabolic state most appropriate for nutrient-rich blood.

**Figure 5:**
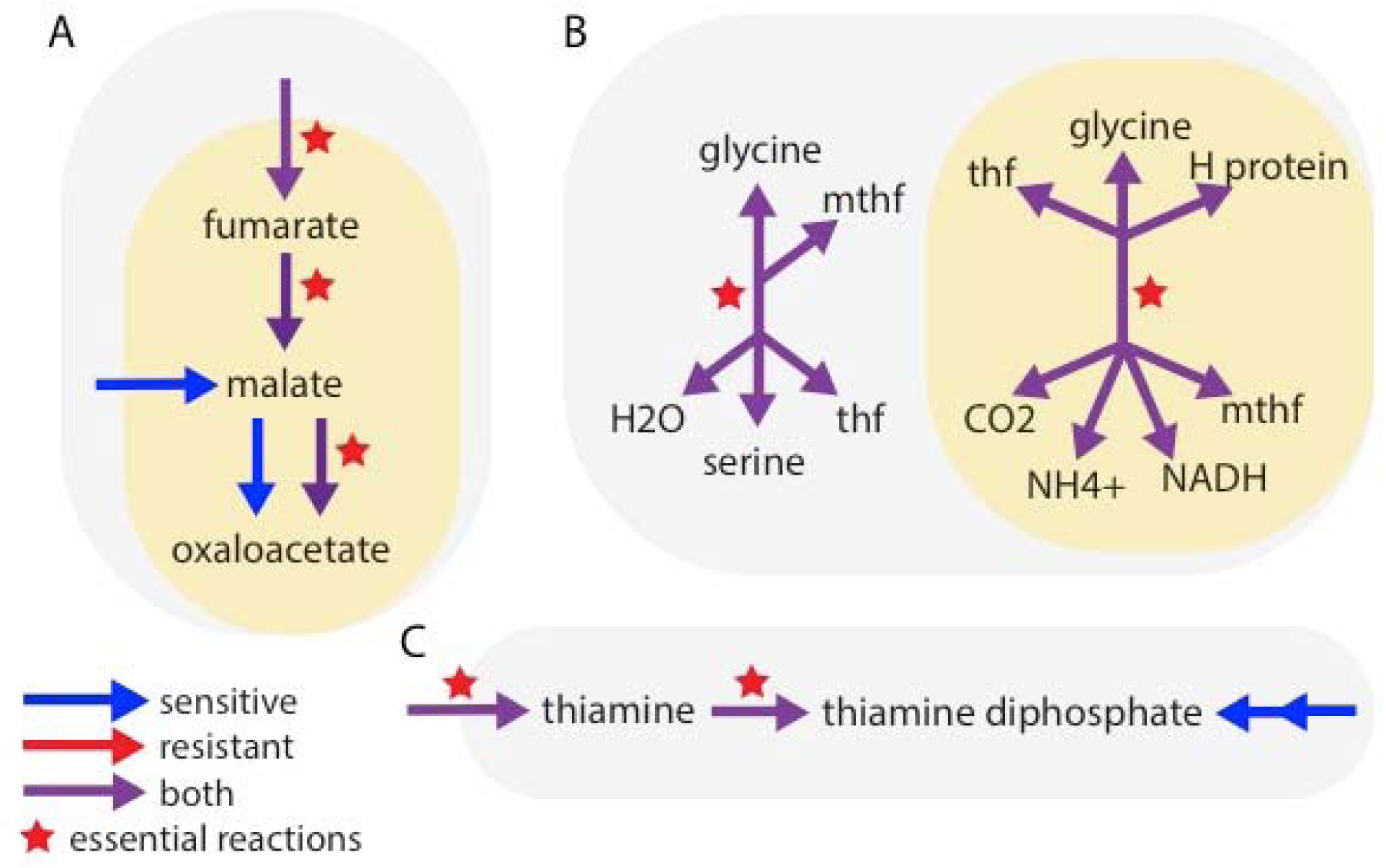
Artemisinin resistance displays unique metabolic weaknesses. ***A* Trycarboxylic acid cycle:** Resistant parasites rely on generation of oxaloacetate from the conversion of fumarate to malate, using fumarate hydratase and malate dehydrogenase, in the mitochondria. Sensitive parasites can also import malate into the mitochondria and use an alternative enzyme (malate:quinone oxidoreductase) to convert malate to oxaloacetate. ***B* Folate metabolism:** Inhibition of the SHMT enzyme (**left**) and the glycine cleavage system (**right**) is lethal in resistant parasites. Sensitive parasites can use either of these enzyme complexes interchangeably to produce mthf and thf. ***C* Cofactor synthesis:** The import of thiamine and the conversion of thiamine to thiamine diphosphate via thiamine thiphosphokinase is essential in resistant parasites. Sensitive parasites can also synthesize thiamine diphosphate *de novo*. Arrows colored for flux via FVA and stars for essentiality. *Gray* background indicates cytosolic localization, *yellow* indicates mitochondrial localization. FVA, flux variability analysis. SHMT, serine hydroxylmethltransferase, mthf, methyltetrahydrofolate, thf, tetrahydrofolate.

Additionally, we identified serine hydroxymethyltransferase (SHMT) and thiamine diphosphokinase as potential drug targets of resistant parasites (Table 5, Figure 5B); see below for discussion of SHMT. Both the import of thiamine and thiamine diphosphokinase are essential only in resistant parasites (Figure 5C), and we predict inhibition of import or enzyme activity would specifically target resistant parasites. These reactions are relatively uncharacterized as the parasite can likely synthesize thiamine diphosphate (vitamin B1) *de novo* [126]. Thus, this approach can generate novel hypotheses and be utilized for the identification of novel drug targets, and, importantly, targets to help prevent the development of resistance.

### Data-driven model implementation highlights knowledge gaps

Although iPfal17 represents our best understanding of intra-erythrocytic *P. falciparum* biochemistry as the most comprehensive reconstruction to date, predictions occasionally contradict published experimental results. These results illuminate experimental complexities and incompletely characterized pathways. For example, our model predicted that cytosolic SHMT is only essential in resistant parasites (Figure 5B left). In sensitive parasites, the essential metabolites can be generated by SHMT or the mitochondrial glycine cleavage system, given the reversible nature of these enzymes [127, 128]. Therefore, in our sensitive models, neither SHMT nor the glycine cleavage system is essential when knocked out individually. This observation conflicts with the literature, as SHMT is essential in cultured parasites [127, 129, 130]. Thus, iPfal17 is unable to predict this intricacy of parasite metabolism, revealing interesting regulatory effects, an uncharacterized location dependency for metabolite generation, or *in vivo/in vitro* differences in enzyme reversibility.

Similarly, model integration reveals that protein localization influences essentiality predictions. We predicted that the cyclical oxidization and reduction of glutathione, a key regulator of oxidative stress [131-134], and supporting reactions were essential only in resistant parasites when the glutathione redox system was located within the mitochondria (data not shown). This is consistent with artemisinin’s induction of reactive oxygen species, the parasite’s obvious need to survive this stress [24, 33-35], and data showing artemisinin sensitivity is correlated with glutathione levels in rodent *Plasmodium* [27]. However, upon moving these reactions to the cytosolic and apicoplast compartments (as supported by [135]), these reactions were no longer essential. Thus, model analysis challenges the integration of previously incomparable datasets by demonstrating that this localization and role of glutathione yield different predictions. Future studies will be required to clarify these findings.

Here, we have presented a novel blood-stage-specific *P. falciparum* metabolic network reconstruction, iPfal17, and investigation of the metabolic differences between artemisinin sensitive and resistant parasites. Antimalarial resistance is a major public health problem and we demonstrate that constraint-based modeling can be used to reveal metabolic shifts that arise with or in support of the resistant phenotype and discrepancies between otherwise incomparable datasets. We find inherent differences in artemisinin resistant and sensitive parasite metabolism, even before artemisinin treatment. Artemisinin resistant parasites have major metabolic shifts in the mitochondria and in the synthesis of folates and polyamines, indicating incomplete transition to the metabolic state most appropriate for the blood-stage environment. These findings generate areas of future research to elucidate *Plasmodium* biochemistry, understand the evolution of artemisinin resistant parasites, and tackle antimalarial resistance.

## METHODS

### Expression analysis

Normalized preprocessed data was obtained from GEO (GSE59097) [49]. Probes on the microarray platform GPL18893 were annotated using NCBI’s stand-alone BLAST correcting the gene labels for 647 probes. Only top hits were used; specifically, hits with greater than 95% identity, no gaps, and a score of over 100 were used (**Suppl. Table 8**). The R package limma was used to compare artemisinin sensitive and resistant samples collected from Cambodia and Vietnam [136]. Samples with predominantly ring-stage parasites with no detectable gametocytes were used. Resistant parasites were defined as both having at least one mutant Kelch13 allele and a parasite clearance half-life of greater than 5 hours (Figure 1) [49, 137]. Sensitive parasites were defined by having at least no mutant Kelch13 alleles and a parasite clearance half-life of less than 5 hours. Random Forest classifiers were built using the R package randomForest, using all ring-stage samples [138]. The metadata classifier used the following variables as outlined in the original study [49]. Cambodian and Vietnamese ring-stage transcriptomes were compared separately to ensure patterns associated with resistance status were reproducible across phylogenies. These countries were chosen for large number of isolates and prevalence of resistance. Microarray probes were screened to remove non-metabolic genes and to keep only one probe per gene (consistent with standard practice). Multiple testing correction was conducted using a false discovery rate [139, 140].

Gene expression data with calculations of fold changes and associated adjusted p-value were incorporated into our curated model using the Metabolic Adjustment for Differential Expression algorithm (MADE). Briefly, MADE statistical significance of gene expression changes along with network context to assign binary gene states (‘on’/‘off’) to each metabolic gene, constraining the network by limiting flux through reactions mapped to ‘off’ genes while maintaining growth, or a similar objective. An 80% growth threshold was used given that there is no reported evidence that resistant and sensitive parasites produce variable biomass as measured by the size of ring-stage parasites; while varying this threshold affects sensitive parasite biomass yield, it does not affect essentiality predictions (data not shown). Essential genes were predicted for the resultant condition-specific models (Figure 3) by conducting single gene and reaction deletions with established algorithms [141]. Consensus lethal gene and reaction deletions from the Cambodian and Vietnamese parasite models were used.

### Flux Analysis and Metabolic Tasks

Flux balance analysis (FBA) is an approach to explore metabolic phenotypes *in silico* [142]. FBA simulates steady-state flux values for each of the network’s reactions that maximize subsequent flux through an objective function given a set of constraints. We chose biomass production as the objective reaction, consistent with previous studies interrogating gene essentiality [50, 53, 96, 143], and permitted flux through all transport reactions. Constraints on the system include conservation of mass, reversibility of reactions, and reaction localization. Flux variability analysis (FVA) uses a related approach to find the range of fluxes permissible given system constraints [144].

We simulated *in vitro* growth requirements by modifying media components or access to particular metabolites. Metabolite import or production was eliminated from the reconstruction, and subsequent biomass production was observed. Lethal modifications were defined as changes that resulted in no production of biomass; growth-reducing modifications were defined as producing less than 90% of unconstrained flux value [98, 143].

### Curation

Manual curation of an existing *P. falciparum* metabolic network reconstruction [50] was conducted by a literature review and reference to generic and *Plasmodium*-specific databases (KEGG, Expasy, and PlasmoDB, MPMP) [43, 145-147]. Data obtained from these sources were used to evaluate the inclusion of reactions as well as their stoichiometry, reversibility, localization, and gene annotations. Genetically and biochemically supported reactions were kept and new reactions were added. Reactions were removed if (1) explicitly determined to be false or (2) were nonfunctional and not supported biochemically or genetically. Spontaneous reactions (reactions that occur without enzymes) are noted to differentiate from orphan reactions (reactions with unknown enzyme catalysts).

In order to assess gene essentiality, we used a biomass reaction as the modeling objective function. Thus, flux through this reaction, simulating cellular growth, was maximized for all *in silico* experimental procedures. We used the biomass reaction from a previous study [50] with modifications. Curation of the biomass reaction was informed by metabolites detected in metabolomics studies [28, 56-58]; if possible, metabolite ratios were predicted from metabolomics data. We curated the biomass reaction with in consideration of published essentiality data; metabolites detected in metabolomics experiments with no known catabolism or import pathways were excluded from the biomass reaction.

### Essentiality studies

We predicted essentiality by performing single deletion studies with both genes and reactions and double gene deletion studies in our curated model and each expression-constrained sensitive and resistant models. Gene deletions were simulated by removing the gene of interest from the model. This change results in the inhibition of flux through all reactions that require that gene to function. If the model could not produce biomass with these constraints, the gene was deemed essential. Growth reducing phenotypes were also observed and noted. For reaction deletion studies, we removed reactions sequentially. Subsequent growth effects were used to determine reaction essentially. Consensus results for resistant or sensitive models are discussed.

## LIST OF ABBREVIATIONS

MADE: Metabolic Adjustment for Differential Expression algorithm
SHMT: serine hydroxymethyltransferase
TCA: tricarboxylic acid
FVA: cycle Flux variability analysis
FBA: Flux balance analysis

## Acknowledgements

We acknowledge the members of the Guler, Papin, and Petri labs for their thoughtful conversations and insight. We would also like to thank Dr. Paul Jensen, Dr. Edik Blais, and Gregory Medlock for project feedback.

## Availability of data and materials

The supporting data and materials of this article are included within the article and additional files; additionally, our model and code is available on https://github.com/maureencarey/iPfal17.

## Authors’ contributions

MAC, JG, and JP designed the study. MAC curated the model and performed metabolic network, statistical, and machine learning analyses. All the authors participated in data interpretation, and read and approved the manuscript.

## Additional files

Additional information is available upon request, in Additional file 1 (Supplemental Figures 1-4), Additional file 2 (Supplemental Tables 1-8), and Additional file 3 (model).

---

**Supplemental Figure 1:**
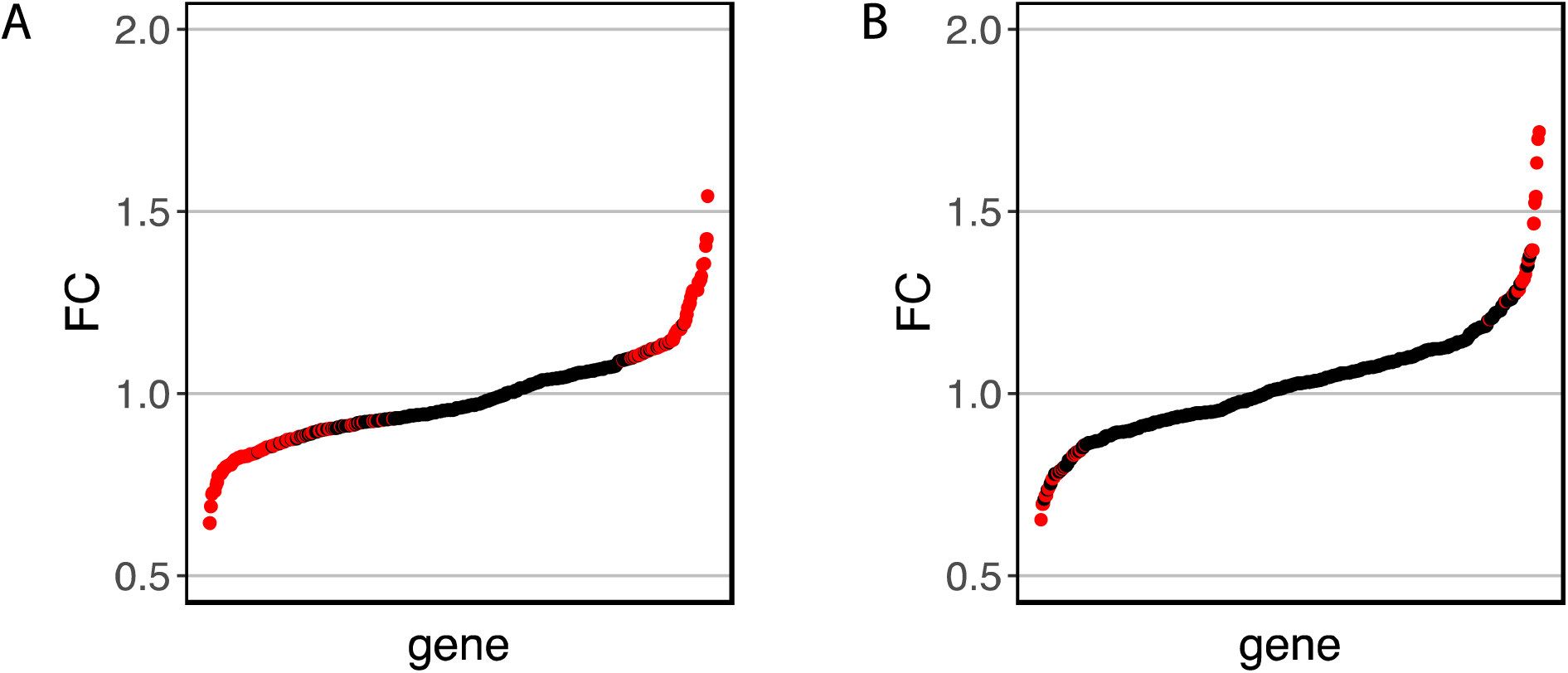
Distribution of genome-wide expression data demonstrates moderate differential expression between sensitive and resistant parasites. Fold change values from differential expression between sensitive and resistant parasites from Cambodia (***A***) and Vietnam (***B***) with significantly differentially expressed genes in *red*.

**Supplemental Figure 2:**
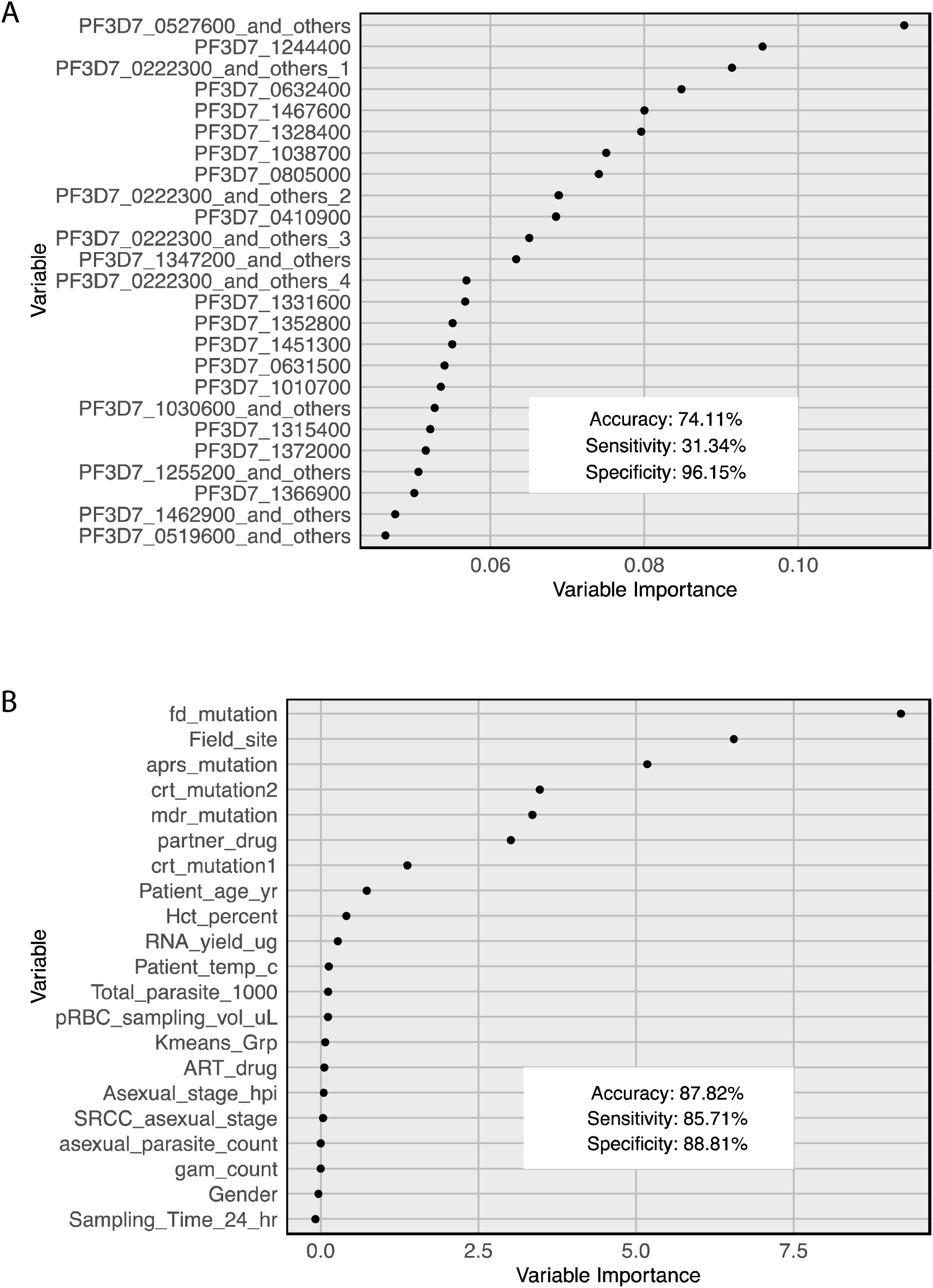
Artemisinin resistance is better predicted by metadata classifier than expression classifier. Using full expression profile (**A**) or metadata (**B**), including patient and parasite features (see Methods), we can classify samples as artemisinin sensitive or resistant by Random forest analysis. Of the top 25 most important variables (gene probes) in the expression classifier, 12 encoded exported proteins, four genes of complete unknown function, three encoded a putative kinase and putative phosphatases, one encoded a component of dynein, four were uncharacterized genes though to be involved in protein folding or trafficking, and one encoded a transcription factor. Abbreviation key: (all from [49] unless noted otherwise) aprs_mutation = apicoplast ribosomal protein S10 (PF3D7_1460900.1) mutation [14];fd_mutation = ferredoxin (PF3D7_1318100) mutation [14];Field_site = location at which blood sample was collected; mdr_mutation = multidrug resistance protein 2 (PF3D7_1447900) mutation;partner_drug = Partner drug (Artemisinin based combination treatment) administered from day 3 onwards; crt_mutation2 = second CRT (PF3D7_0709000)[14] mutation measured; crt_mutation1 = first CRT (PF3D7_0709000)[14] mutation measured; Patient_age_yr = patient age in years; pRBC_sampling_vol_uL = Volume of packed RBC collected (uL); RNA_yield_ug = Amount of Total RNA isolated for each sample (μg)□; Patient_temp_c = patient temperature at time of admission in Celsius; ART_drug = Type and dosage of artemisinin drug given once a day on days 0, 1 and 2; asexual_parasite_count = Total asexual parasite densities per uL on admission; total_parasite_1000 = total number of parasites in whole sample of infected RBC collected (pRBC collection vol. * total parasite count per uL) divided by 1000□; SRCC_asexual_stage = Spearman rank correlation coefficient of the gene expression for the isolate sample to the projected hpi; Kmeans_Grp = expression group (see [49]); Asexual_stage_hpi = Projected hours post invasion (hpi) of the parasite asexual stage; Gender = Patient gender; gam_count = Total gametocyte parasite densities per uL on admission; Hct_percent = patient hematocrit (%) on admission □; Sampling_Time_24_hr = Time of sample collection in 24 hour format □

**Suppl. Figure 3:**
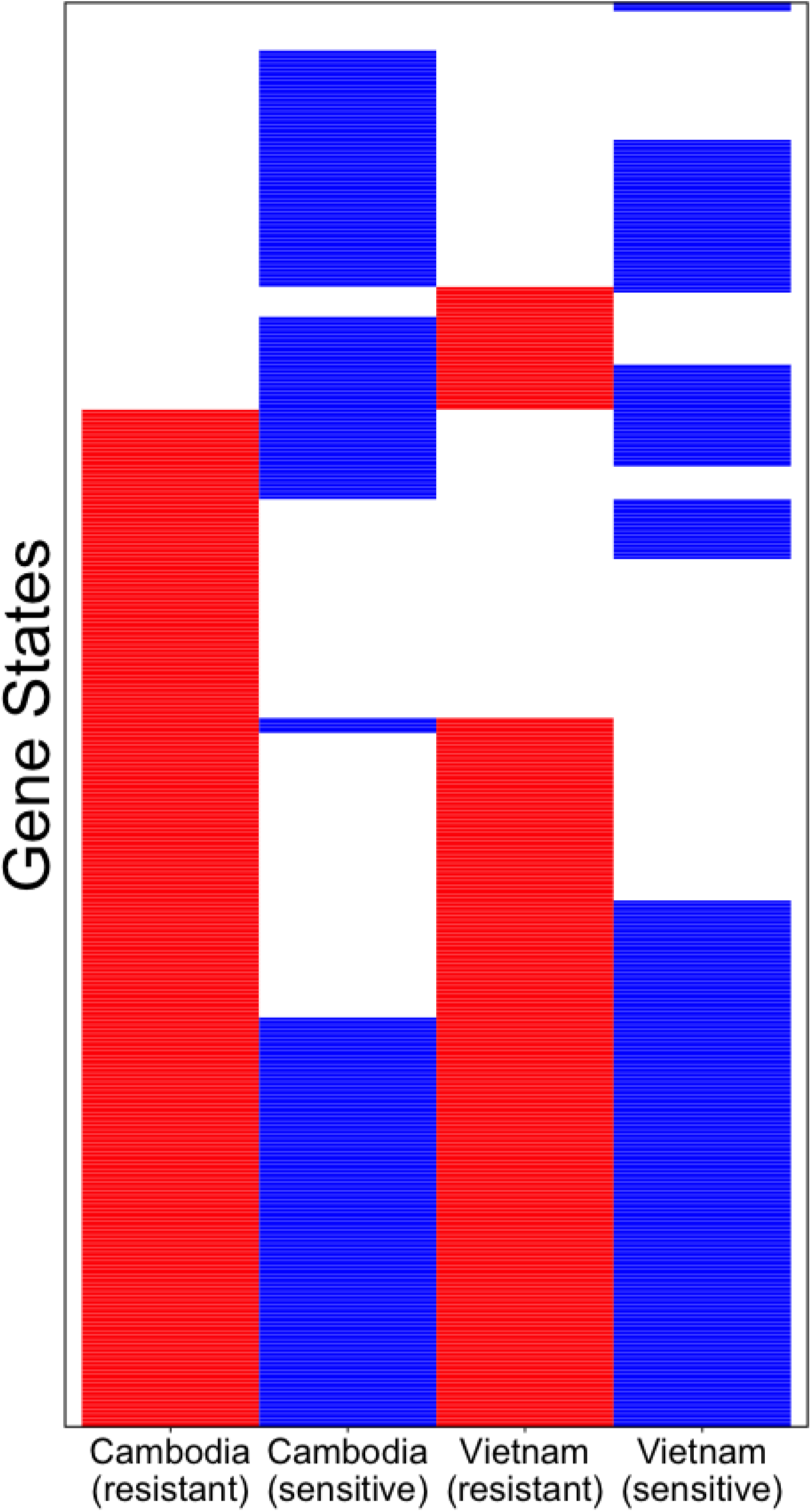
Functional differences in data-driven sensitive and resistant models. Gene states from four condition-specific models, the results of MADE integration, cluster by sensitivity not by location. Active genes in *red/blue*, with genes removed from expression-constrained models in *white*.

**Suppl. Figure 4:**
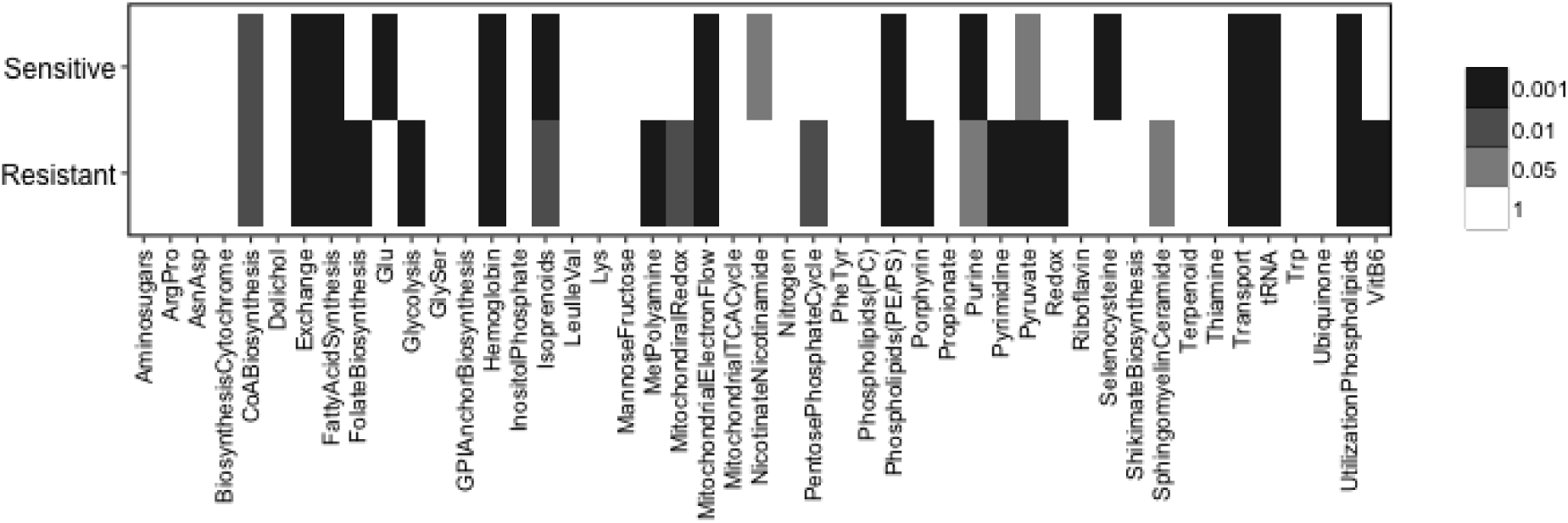
Artemisinin sensitive and resistant parasites utilize different metabolic genes and pathways. Enrichment analysis of gene utilization in sensitive and resistant parasite models demonstrates functional differences in expression data integration. Consensus gene utilization from resistant and sensitive models (both Cambodian and Vietnamese datasets) were used and compared to unconstrained model. Black = p < 0.001, grey = p < 0.01, light grey = p < 0.05, white = non significant. Abbreviation key: Aminosugars = amino sugar metabolism; ArgPro = arginine and proline metabolism; AsnAsp = asparagine and aspartate metabolism; BiosynthesisCytochrome = biosynthesis of cytochromes; CoABiosynthesis = coenzyme-A biosynthesis; Dolichol = dolichol metabolism; Exchange = exchange reactions; FattyAcidSynthesis = fatty acid synthesis; FolateBiosynthesis = folate biosynthesis; Glu = glutamate metabolism; Glycolysis = glycolysis; GlySer = glycine and serine metabolism; GPIAnchorBiosynthesis = GPI anchor biosynthesis; Hemoglobin = hemoglobin degradation (including hemozoin formation); InositolPhosphate = inositol phosphate metabolism; Isoprenoids = isoprenoid metabolism; LeuIleVal = leucine, isoleucine, and valine metabolism; Lys = lysine metabolism; MannoseFructose = mannose and fructose metabolism; MetPolyamine = methionine and polyamine metabolism; MitochondrialElectronFlow = mitochondrial electron transport chain; MitochondrialTCACycle = mitochondrial tricarboxylic acid cycle; NicotinateNicotinamide = nicotinate and nicotinamide metabolism; Nitrogen = nitrogen metabolism; PentosePhosphateCycle = pentose phosphate cycle; PheTyr = phenylalanine and tyrosine metabolism; Phosphatidylcholine = phosphatidylcholine metabolism; PhosphatidyletanolaminePhosphatidylserine = phosphatidyletanolamine and phosphatidylserine metabolism; Porphyrin = porphyrin metabolism; Propionate = propionate metabolism; Purine = purine metabolism; Pyrimidine = pyrimidine metabolism; Pyruvate = pyruvate metabolism; Redox = redox metabolism; RedoxMitochondrialAntioxidantSystem = mitochondrial redox metabolism; Riboflavin = riboflavin (vitamin B2) metabolism; Selenocysteine = selenocysteine metabolism; ShikimateBiosynthesis = shikimate biosynthesis; SphingomyelinCeramide = sphingomyelin and ceramide metabolism; Terpenoid = terpenoid metabolism; Thiamine = thiamine biosynthesis; Transport = transport reactions; tRNA = tRNA and protein synthesis; Trp = tryptophan metabolism; Ubiquinone = ubiquinone metabolism; UtilizationPhospholipids = utilization of phospholipids; VitB6 = pyridoxal (vitamin B6) metabolism

Suppl. Table 1: Modifications and reaction additions in iPfal17 curation.

Suppl. Table 2: Reactions deleted from Plata *et al.* model in generating iPfal17.

Suppl. Table 3: Predicted lethal reactions in wild-type blood-stage *Plasmodium falciparum*.

Suppl. Table 4: Predicted lethal genes in wild-type blood-stage *Plasmodium falciparum*.

Suppl. Table 5: Consensus predicted lethal reactions across 4 expression-constrained models.

Suppl. Table 6: PlasmoGem orthologous results.

Suppl. Table 7: Extended table 4

Suppl. Table 8: Microarray platform blast results.

